# PURE-seq identifies *Egr1* as a Potential Master Regulator in Murine Aging by Sequencing Long-Term Hematopoietic Stem Cells

**DOI:** 10.1101/2024.08.12.607664

**Authors:** Sixuan Pan, Kai-Chun Chang, Inés Fernández-Maestre, Stéphane Van Haver, Matthew G. Wereski, Robert L. Bowman, Ross L. Levine, Adam R. Abate

## Abstract

Single-cell transcriptomics is valuable for uncovering individual cell properties, particularly in highly heterogeneous systems. However, this technique often results in the analysis of many well-characterized cells, increasing costs and diluting rare cell populations. To address this, we developed PURE-seq (PIP-seq for Rare-cell Enrichment and Sequencing) for scalable sequencing of rare cells. PURE-seq allows direct cell loading from FACS into PIP-seq reactions, minimizing handling and reducing cell loss. PURE-seq reliably captures rare cells, with 60 minutes of sorting capturing tens of cells at a rarity of 1 in 1,000,000. Using PURE-seq, we investigated murine long-term hematopoietic stem cells and their transcriptomes in the context of hematopoietic aging, identifying *Egr1* as a potential master regulator of hematopoiesis in the aging context. PURE-seq offers an accessible and reliable method for isolating and sequencing cells that are currently too rare to capture successfully with existing methods.

## Introduction

Single-cell transcriptomics is powerful for elucidating the properties of individual cells and can discover phenotypes without relying on predetermined genes or markers. This makes it useful in highly heterogeneous systems with unknown cell properties^1–4^. However, its unbiased nature often leads to the analysis of abundant, well-characterized cellular states at the expense of rare cell populations and increased cost^5,6^. An enrichment method that selectively captures rare cell populations while removing unwanted cells can increase the coverage of rare cells, enabling deeper analysis at the same cost.

Several methods exist for enriching target cells before single-cell sequencing, typically using antibody-based capture approaches to label and isolate cells of interest. Techniques such as fluorescence-activated cell sorting (FACS), magnetic-activated cell sorting (MACS), and cell levitation isolate cells based on expression of specific surface markers^7–9^. However, current single-cell methods do not directly integrate with the output of a flow cytometer, necessitating a transfer step that can result in cell loss or degradation, compromising data quality. This is especially problematic for extremely rare cell applications where the number of captured cells may be too low for reliable transfer. Other alternatives, such as direct cell sorting into well plates or using nanowell array chips, involve labor-intensive workflows and have limited throughput capabilities^10,11^. An ideal approach would allow the flow cytometer to directly load cells into the high-throughput single-cell RNA-sequencing (scRNA-seq) apparatus, minimizing handling, ensuring the highest data quality, and capturing rare cell populations; however, this is not possible with existing methods.

In this paper, we introduce PURE-seq (PIP-seq for Rare-cell Enrichment and Sequencing), a method for sequencing rare cells. PURE-seq is based on our development, Pre-templated Instant Partition sequencing (PIP-seq) ^12^, which allows scalable scRNA-seq without microfluidics using a fully encapsulated Eppendorf tube. The compact nature of the PIP-seq reservoir and its compatibility with standard Eppendorf tubes, commonly used in flow cytometry, enable direct cell loading from the flow cytometer into the PURE-seq reaction. This eliminates additional handling, reducing cell loss and degradation. The tube is vortexed immediately after cell loading, encapsulating, and lysing the cells in droplets within one minute for the PIP-seq single-cell barcoding workflow^12^. This simplicity and minimal handling allow reliable capture of extremely rare cells; 60 minutes of sorting can capture tens of cells at a rarity of 1 in 1,000,000. The rarity of cells captured scales with sorting duration, allowing even rarer cells to be sequenced with more sorting time.

Using PURE-seq, we analyzed murine long-term hematopoietic stem cells (LT-HSCs), a rare and heterogeneous bone marrow (BM) population difficult to recover in sufficient numbers for high-quality scRNA-seq with current methods^13,14^. PURE-seq enabled us to characterize their transcriptomes in low-input samples. Previous studies hinted at the role of *EGR1* in human LT- HSCs^15,16^, but its exact function in mice is unclear. These studies demonstrate higher *EGR1* expression in aged human hematopoietic stem and progenitor cells (HSPCs), suggesting EGR1 may regulate quiescence, proliferation, and localization. Attenuated expression of EGR1 might decrease senescence and activate aged HSPCs, offering a potential target for rejuvenation strategies^17^. PURE-seq allowed us to recover sufficient cell numbers to identify *Egr1* as a potential master regulator gene in the aging of murine LT-HSCs. Here, we show that PURE-seq provides a simple workflow to sort and sequence rare cell populations, which is arduous with existing methods, and reliably recapitulates data generated by standard 10x Genomics.

## Results

### PURE-seq: Direct FACS sorting of target cells into PIP-seq single-cell RNAseq reactions

The PURE-seq workflow utilizes readily available commercial platforms, FACS and PIP-seq, to achieve scalable, reliable, and accessible sequencing of extremely rare cells. In PURE-seq, cells are sorted directly into single-cell barcoding reaction tubes. Subsequent cell encapsulation follows the standard PIP-seq protocol^12^, which involves adding encapsulation oil, vortexing for one minute, lysing cells, and capturing mRNA (**Figure 1**). To optimize cell viability and capture efficiency, we fine-tuned cell sort stream alignment, sorting speed, and total sorting duration (**Methods**).

**Figure 1.**
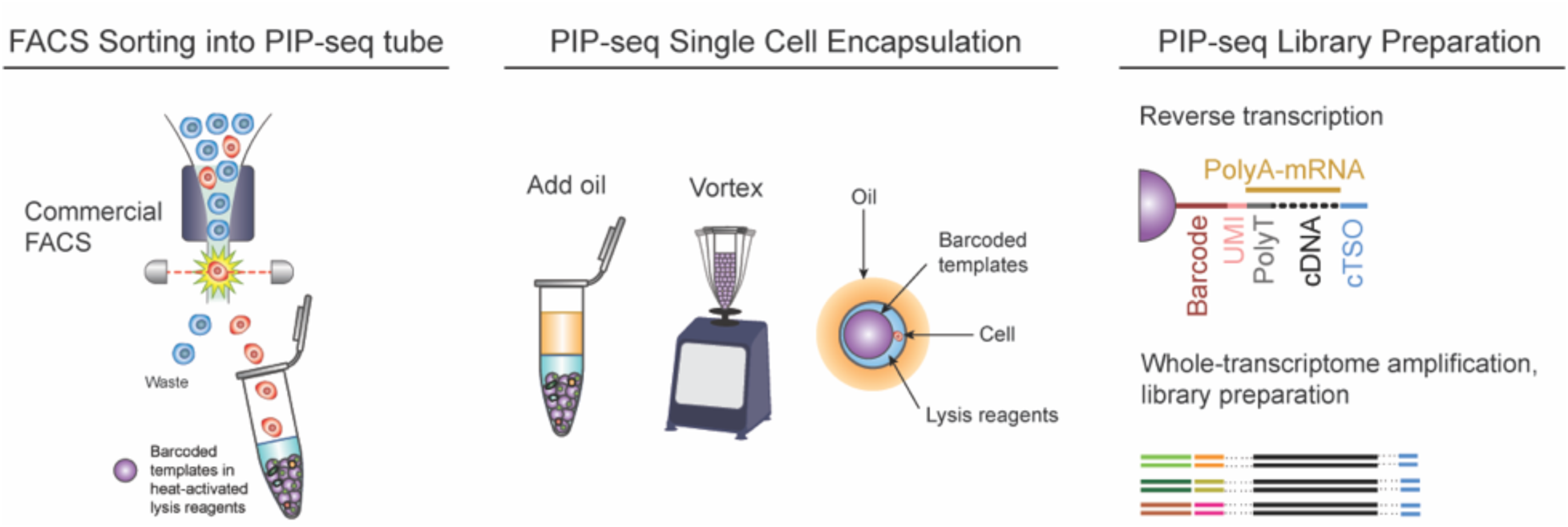
Workflow of PURE-seq with enriched and sorted rare cells from a heterogeneous population. PURE-seq utilizes a commercial FACS system to sort fluorescently labeled target cells directly into PIP-seq reaction tubes containing barcoded templates in heat-activated lysis reagents. The subsequent single-cell encapsulation in droplets follows the standard PIP-seq protocol^12^, which involves adding oil, vortexing, heat-activated lysis, and capturing mRNA on the barcoded templates. After mRNA capture, reverse transcription, and whole-transcriptome amplification are conducted in bulk to prepare barcoded cDNA for Illumina sequencing.

Fluorescence-activated cell sorters have multiple sorting precision modes. In “single-cell” mode, sorting specificity is prioritized, and ambiguous results due to staining, cell clumping, or coincidences in the detector are discarded. In “yield” mode, ambiguous events are recovered to ensure maximum retrieval of rare cells, even at the cost of capturing some off-target cells. With PURE-seq, we can prioritize capturing rare cells over the purity of the sorted population, leveraging the high single-cell sequencing capacity downstream. For example, PIP-seq reactions can be scaled to accommodate inputs of 2,000, 20,000, and over 100,000 cells^12^. This high capacity is especially useful for sequencing extremely rare cell populations, allowing us to maximize the capture of rare cells during the flow cytometry step. While the final single-cell sequenced population may contain off-target cells, the overall enrichment from pre-sort to post-sort is significant.

To assess the efficacy of PURE-seq, we conducted a human-mouse species-mixing experiment, introducing human HEK 293T cells into mouse NIH 3T3 cells at a dilution of 1 in 1,000. The human (HEK 293T) cells served as the representative target cells within a background population. We labeled the human and mouse cells with different Calcein dyes (**Methods**) and processed the sample using the BD FACS Aria system. We instructed the instrument to sort the first 2,500 human cells into the PIP-seq reaction. In parallel, we used PIP-seq to sequence the unsorted population. For the unsorted population, we recovered no human HEK 293T cells since the rarity was 1 in 1,000, and sequencing just 2,500 cells resulted in no random capture of human cells. By contrast, in the sorted reaction, we recovered 584 human (HEK 293T) cells and 112 off-target mouse (NIH 3T3) cells, illustrating significant enrichment for the target population (**Figure 2A**).

**Figure 2.**
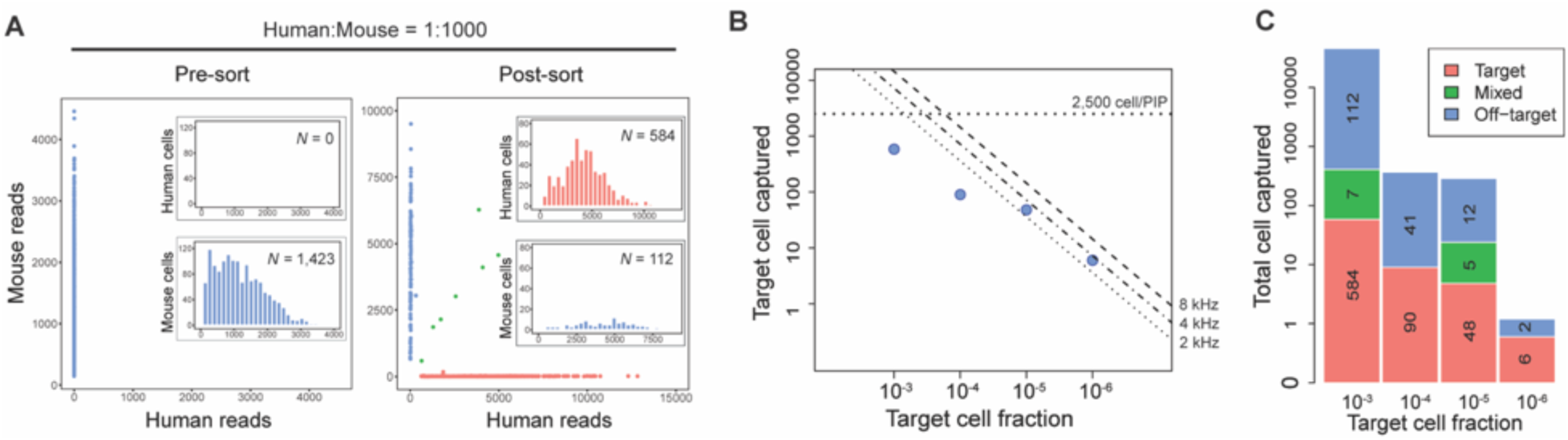
PURE-seq efficiently captures and sequences rare cells isolated by FACS. A) Barnyard plots of mixed human-mouse population (Human:Mouse = 1:1000) sequenced before (left) and after sorting (right). Inserts are histograms of read distribution for sequenced human or mouse cells. Cells are colored by cell type (blue, mouse reads; red, human reads; green, mixed reads). **B)** Number of target cells captured as a function of target cell fraction. The dashed lines mark the theoretical limit of the captured cells. A maximum of 2,500 cells are sorted into each T2 PIP-seq reaction. Contour lines are the theoretical numbers of target cells that can be sorted within 60 minutes with different sorting rates (8 kHz, 4 kHz, and 2 kHz). Blue dots are the actual number of cells sequenced for the mixed human-mouse population with target cell fractions of 10^-3^, 10^-4^, 10^-5^, and 10^-6^. **C)** Number of target and off-target (mis-sort) cells sequenced for each rarity group.

To confirm successful scRNA-seq, we generated barnyard plots, plotting the number of mouse reads for each cell versus the number of human reads it contains. The two populations aligned along the axes, illustrating that most captured cells had either pure mouse or human transcriptomes. We observed some mixed transcriptomes along the diagonal, consistent with co-encapsulation of mouse and human cells during the PIP-seq barcoding step, as is typical in single-cell reactions relying on limiting dilution. These results demonstrate that PURE-seq allows reliable single-cell sequencing of the target cell population for the spiked population at the 1:1,000 rarity level.

A major strength of flow cytometry is its capacity for high-throughput cell sorting, allowing the screening of vast populations to identify rare cellular states. In this experiment, we sought to determine the maximum rarity compatible with PURE-seq. Therefore, we tuned sorting parameters to maximize the total number of cells that could be sorted while minimizing the impact on the cells. We set a maximum sorting duration of 60 minutes and speed of 8 kHz to prevent perturbation of gene expression due to long waiting times and high shear forces in the sorter, respectively, allowing 28.8 million cells to be sorted per run. At peak efficiency, this setup can, therefore, recover cells with a rarity of approximately 1 in 1 million, delivering tens of target cells to the PIP- seq reaction. Thus, the sequencing reaction must be exceptionally efficient to reliably barcode such a tiny number of inputs; typical cell inputs for commercial single-cell instruments exceed 1,000 cells per reaction. Since the maximum input volume for the PIP-seq T2 kit is 5 μL, we also restricted the maximum number of sorted cells to 2,500 based on the droplet volume of the BD FACS Aria system (1.81 nL/drop). If more sorted cells are desired, multiple PIP-seq T2 tubes can be used, or larger PIP-seq kits, such as the T20 (20,000 cells) and T100 (100,000 cells)^12^, can be utilized instead.

With the abovementioned parameters, we assessed the limits of enrichment possible with PURE- seq by conducting sorting experiments at different target cell rarity (**Figure 2B, Supplementary Figure 1**). We confirmed that for target cell rarity ranging from 10^-3^ to 10^-6^, between 564 and 6 target cells, respectively, could be captured and sequenced with 75% or greater purity (**Figure 2C**). This purity level can be increased to 98% by switching from “yield” sorting precision mode to “single-cell” mode, although this reduces the number of recovered cells by ∼33% (**Supplementary Figure 2**).

### PURE-seq significantly increases the capture of LT-HSCs compared to the pre-sort control

LT-HSCs are a rare population in the mouse BM and lie at the top of the hematopoietic hierarchy^18^. Profiling LT-HSCs in scRNA-seq studies has been especially challenging due to their rarity and heterogeneity, which makes it difficult to capture enough true LT-HSCs for detailed analysis^13,14^. To demonstrate the utility of PURE-seq for the analysis of primary samples, we used it to investigate murine LT-HSCs sorted from Lineage^−^Sca-1^+^c-Kit^+^ (LSK) cells based on the expression of SLAM markers, which enrich for HSCs (CD150^+^CD48^−^ LSK cells)^19^. Specifically, to demonstrate how PURE-seq can increase the capture of LT-HSCs compared to a pre-sort control and provide a high-quality dataset to gain biological insights, we studied LT-HSCs throughout murine aging. We harvested whole BM cells from young (2-3 months old), middle-aged (12-14 months old), and old (18-20 months old) C57BL/6 mice. We removed lineage-positive cells to enrich for hematopoietic stem/progenitor cells (HSPCs) before starting the PURE-seq workflow, which encompassed LT-HSC sorting from BM pools (n=2-3 mice/pool) followed by the PIP-seq pipeline and Illumina sequencing (**Figure 3A**). After processing and SCT-transforming the samples with Seurat v4, our analysis revealed that 19.37% expressed both *Sca-1* and *c-Kit* and that 7.27% could be considered LT-HSCs by including the expression of *Slamf1*, which encodes for the phenotypic cell surface marker CD150^20^ (**Figure 3B**).

**Figure 3.**
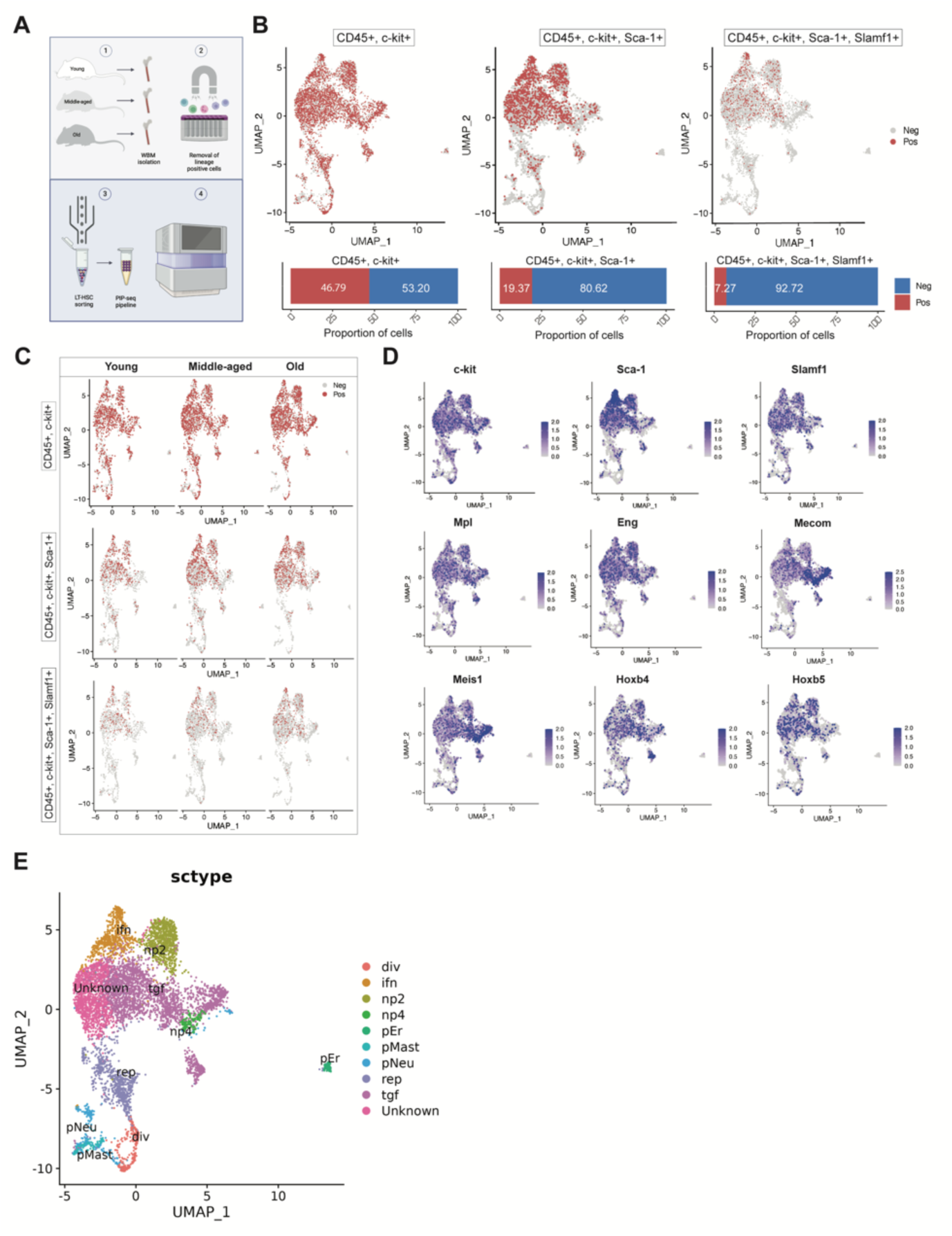
PURE-Seq isolates murine long-term repopulating hematopoietic stem cells and enables single-cell sequencing via PIP-seq and analysis throughout aging. A) Schematic of the PURE-seq pipeline for sorting murine LT-HSCs from young, middle-aged, and old mice after depleting lineage-positive cells for scRNA-seq library preparation using PIP-seq and Illumina sequencing. **B)** Comparison of hematopoietic cells (CD45^+^) expressing *c-Kit* only; *c-Kit* and *Sca-1*; or *c-Kit*, *Sca-1*, and *Slamf1*, simultaneously in the integrated UMAP plot from the dataset (top) and breakdown bar graphs of the total percentages of positive and negative cells (bottom) **C)** UMAP plots showing differences in the numbers of *c- kit* only; *c-Kit* and *Sca-1*-double positive; or *c-Kit*, *Sca-1*, and *Slamf1*-triple positive cells across murine aging. **D)** UMAP plots from the integrated dataset showing cells expressing key LT-HSC signature genes. **E)** UMAP displaying identified cell populations in the integrated dataset annotated according to Hérault *et al.* ^21^

We observed that LT-HSCs did not express CD48, consistent with our FACS gating strategy, which excluded CD48^+^ cells (**Supplementary Figure 3**A). Our analysis also revealed that the percentage of LT-HSCs increased with age (**Supplementary Figures 3A, B**), which aligns with previous studies demonstrating an increase in their percentage within the aged BM^22,23^. This was further confirmed by the generation of Uniform Manifold Approximation and Projection (UMAP) plots that showed a higher number of hematopoietic cells expressing *Kit*, *Sca-1*, and *Slamf1* genes in the middle-aged and old samples compared to their young counterparts (**Figure 3C**). *Kit*^+^, *Sca- 1*^+^, *Slamf1*^+^ cells clustered in the head region of the UMAP plot, co-localizing with the expression of key LT-HSC genes such as myeloproliferative leukemia virus oncogene (*Mpl*), endoglin (*Eng*), MDS1 (*Mecom*), Meis homeobox 1 (*Meis1*), and homeobox genes (*Hoxb4* and *Hoxb5*) (**Figure 3D**).

As a control, we sequenced pre-sort samples using the PIP-seq pipeline and found that only 0.78% of the cells co-expressed *Kit*, *Sca-1,* and *Slamf1*, indicating that with PURE-seq, we were able to increase the percentage of LT-HSCs by 9.3-fold. Regarding the pre-sort control, we also detected that even though the samples were enriched for HSPCs, there were still differentiated immune cells and non-hematopoietic BM cell types, such as endothelial cells and fibroblasts (**Supplementary Figure 3**C), which highlights the inefficiency of cell enrichment methods, such as MACS for lineage-positive hematopoietic cell depletion (as we used in our experiment). In terms of the post-sort samples, 6,725 cells that passed the Seurat quality control were captured, with an average of 841 cells per sample after sorting 2,500 cells with the single-cell mode (**Supplementary Figure 3**D). This demonstrates that 33.64% of the sorted cells were of high quality, a percentage that can be increased using the yield mode, as shown in our sorting precision modes experiment (**Figure 2, Supplementary Figure 2**).

After integrating all the samples, we identified 12 clusters based on transcriptomic differences (**Supplementary Figure 3**D). Next, we used a publicly available dataset from Héuralt *et al.*^21^ to compare their signatures with ours (**Supplementary Table 1**). Similarly, they analyzed LT-HSCs from pooled FACS-sorted LT-HSC samples of old and young mice after the removal of lineage-positive cells, using 10x Genomics instead. They characterized their cell clusters based on differential gene expression analysis in combination with gene set enrichment analysis and gene signatures related to hematopoiesis. Based on their gene markers, we were able to identify 9 out of their 15 cell types, mostly coinciding with non-primed clusters, thus classified because of their lack of expression of lineage-restricted genes (i.e., interferon response (ifn), non-primed (np)2, growth factor signaling (tgf), np4, replicative (rep), and dividing (div)). These non-primed clusters were in the head of the UMAP plot, except for an unknown cluster that did not match any of their signatures, possibly due to the lack of the middle-aged group or other experimental variations in their dataset. We also detected three lineage-primed clusters that were enriched for cells with neutrophil (pNeu) and mastocyte (pMast) or erythroid (pEr) commitment gene markers, but these were located either at the very end of the tail (pMast and pNeu) or clustered completely separately from the bulk of cells (pEr) (**Figure 3E**).

Our dataset was largely comparable to datasets generated with 10X Genomics Chromium, with a predominance of non-primed hematopoietic cell clusters^21^. Furthermore, the good quality metrics across our 12 identified clusters (**Supplementary Figure 3**F), the clear split by biological condition (i.e., age group) with concomitant detection of differences in cell numbers across clusters in our integrated dataset (**Supplementary Figure 3**G), indicated the suitability of PURE-seq as a reliable alternative pipeline to isolate a rare cell population and analyze their single-cell transcriptomes to study their heterogeneity in complex biological phenomena such as hematopoietic aging.

### Subsetting LT-HSCs from the bulk sample allows for analysis of age-related cell cycle and transcriptomic differences

Next, we evaluated the purity of LT-HSCs in our data using the scGate package^24^ (**Supplementary Table 2**). We confirmed that LT-HSCs were indeed dispersed throughout the UMAP plot, with the highest concentration in the head and middle regions of the tail (**Figure 4A**). This aligned with previous findings using Seurat (**Figures 3B, C**). Notably, the distinct cluster that stood apart did not contain any LT-HSCs. Additionally, the end of the tail of the central projection had minimal LT-HSC numbers, which was consistent with the Héuralt *et al*. integration that revealed erythroid, neutrophil, and mastocyte commitment gene expression in these clusters^21^, suggesting that they likely consisted of committed progenitors or were possibly contaminated with differentiated cells.

**Figure 4.**
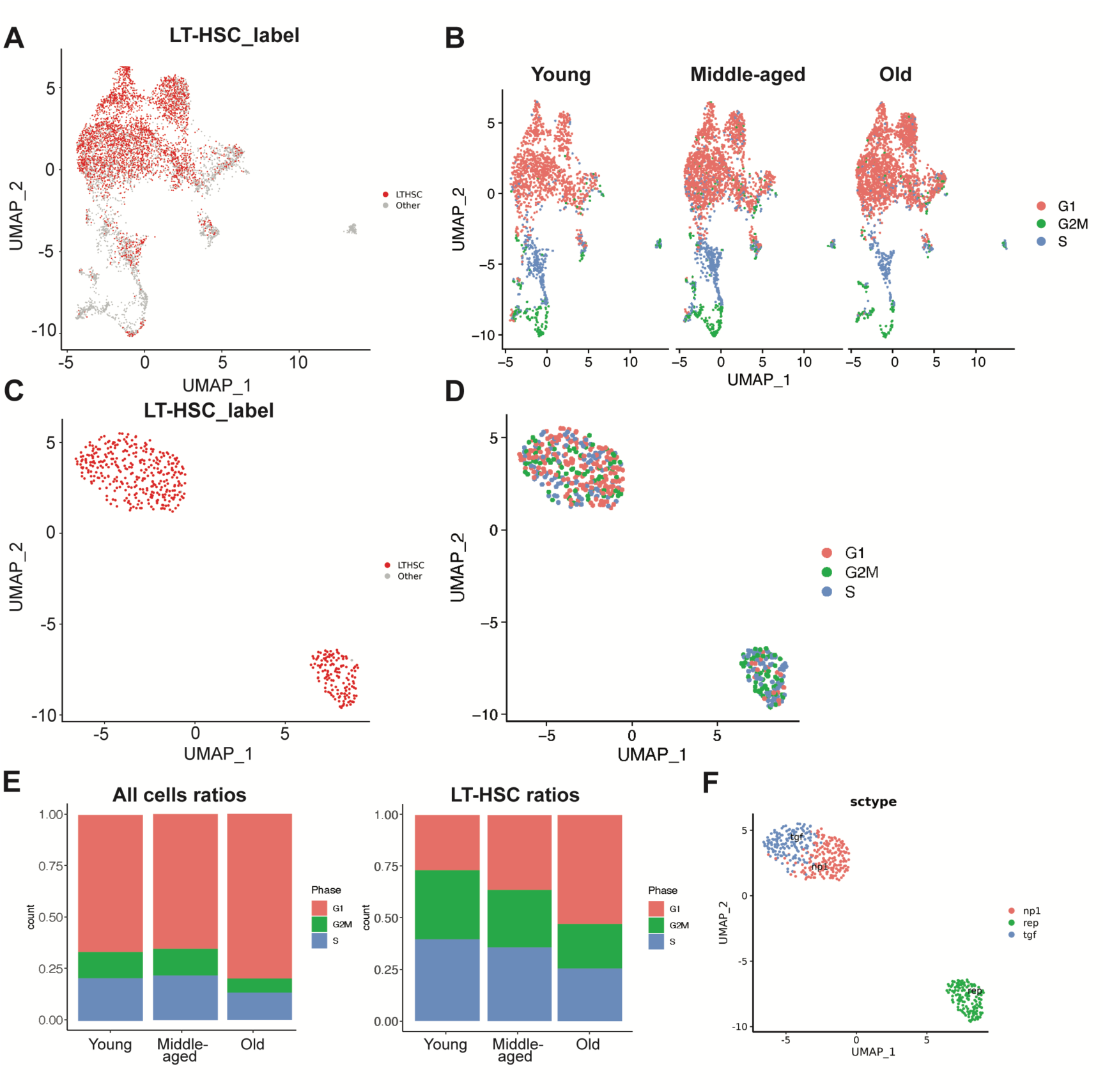
scGATE marker-based purification, cell cycle analysis, and re-clustering of LT-HSCs. A) UMAP plot indicating the purity of LT-HSCs using scGate. **B)** Analysis of cell cycle phases in the integrated UMAP plot. **C)** UMAP plot of re-clustered LT-HSCs as per the scGate label. **D)** Analysis of cell cycle phases in the re-clustered (purified) LT-HSC population. **E)** Stacked bar graphs showing the ratios of all cells (left) or LT-HSCs (right) in different phases of the cell cycle. **F)** UMAP plot of LT-HSCs labeled by cell types as annotated by Hérault *et al.* ^21^

To further validate our dataset, we set out to determine whether age-related cell cycle differences could be detected across the UMAP plot, as such changes are expected with hematopoietic aging. Using the Seurat pipeline, we found that most of the LT-HSC signature overlapped with the G1 phase classification and that the number of cells at the G1 phase appeared to increase with aging (**Figure 4B**). To further examine these differences in LT-HSCs, we extracted the pure LT-HSC population for re-embedding and re-clustering. We identified three distinct clusters (**Supplementary Figure 4**A) where nearly 100% of the cells were labeled LT-HSCs (**Figure 4C**, **Supplementary Figure 4**B). After successfully running a second post-clustering quality control check (**Supplementary Figure 4**C), we observed that G1 phase cells dominated the top clusters (clusters #0 and #1 in **Supplementary Figure 4**A), while cells at G2/M and S phases appeared to preferentially locate within the bottom cluster (cluster #2 in **Supplementary Figure 4**A) (**Figure 4D**). As we observed in the overall integrated sample before extracting the LT-HSCs subset, the proportion of LT-HSCs at the G1 phase increased at the expense of the G2/M and S phases, showing a more significant trend throughout aging compared to that of the larger dataset (**Figure 4E**). We then analyzed the gene expression signatures provided by Héuralt *et al*.^21^, focusing on the LT-HSC subset. We observed that these corresponded to non-primed gene expression, specifically tgf, np1, and rep (**Figure 4F**). The rep signature, characterized by DNA repair and replication genes, had the highest number of cells at the G2/M and S phases, coinciding with cluster #2 (**Supplementary Figure 4**A). These findings support the notion that, despite an increase in their numbers, LT-HSCs have a gradual loss of self-renewal with aging, which has been extensively reported^25^.

Although refining the dataset was possible by extracting and re-clustering the LT-HSC transcriptomes, the use of the overall integrated sample showed enough LT-HSC purity to conduct a representative analysis, as shown by the scGate LT-HSC label (**Figure 4A**), and the expression of relevant LT-HSC genes (**Figure 3D**), as well as markers of undifferentiated HSPCs (e.g., *Procr*) and regeneration/myelosuppression following injury (e.g., *Notch2*), in combination with the nonexistent or low expression of lineage-specific genes, such as the lymphoid-associated interleukin 7 receptor (*Il7r*) and CD79A antigen (*Cd79a*), which drive differentiation towards T/B lymphoid cell lineages (**Supplementary Figure 4**). Additionally, both the overall integrated dataset and the LT-HSC subset allowed for the detection of age-related cell number differences across all the Seurat clusters (**Supplementary Figures 4E, F**). The cross-comparison with the Héuralt *et al.* dataset ^21^ demonstrates that the PURE-seq pipeline can obtain similar results while analyzing over half the number of cells (6,725 versus 15,000 cells) while allowing for the inclusion of an extra condition (the middle-aged group); this ability is especially valuable in sample scarcity scenarios where cell numbers are limiting.

### Identification of EGR1 as a transcription factor determining LT-HSC gene upregulation during aging

Aging causes genetic and epigenetic changes that lead to a decline in HSPC function and self- renewal^26^. Recent studies have identified genes that may regulate hematopoietic aging, revealing differences in gene expression and aging biomarkers, as well as an inclination towards myeloid-biased hematopoiesis as early as middle-age in mice^27,28^. In this context, single-cell transcriptomics has been useful in identifying crucial genes that could be targeted in potential hematopoietic rejuvenation strategies. To explore whether we could identify a relevant gene determining LT-HSC gene upregulation in aging from our dataset, we performed differential gene expression analysis and generated a bubble plot with top-downregulated or upregulated genes during LT-HSC aging (**Figure 5A**). Although most differences laid in the expression of genes involved in fundamental cellular processes, including DNA synthesis (e.g., *Rrm2b*), autophagy (e.g., *Vmp1*), and transcription (e.g., *Cnot6*), we observed that there was an overall elevated expression of genes regulating the immune system and inflammatory responses with aging, as previously shown^27,29^. These genes included jun B proto-oncogene (*Junb*), suppressor of cytokine signaling 3 (*Socs3*), metallothionein (*Mt1*), immediate early response 2 (*Ier2*), Krüppel-like transcription factor 4 (*Klf4*), death-associated protein kinase 1 (*Dapk1*) and genes encoding for members of the S100 protein family (e.g., *S100a6*, *S100a9*). We also found that metabolic genes showed noteworthy differences, including the upregulation of genes implicated in lipid metabolism (e.g., *Slc22a27*), glycogenesis (e.g., *Phkg1*), and growth factor signaling, such as the early growth receptor 2 (*Egr2*) and 3 (*Egr2*), and the expression of *Egr1*, Insulin growth factor 1 receptor (*Igf1r*) and transforming growth factor, beta receptor I (*Tgfbr1*); interestingly, with the latter three peaking in middle age (**Figure 5A**).

**Figure 5.**
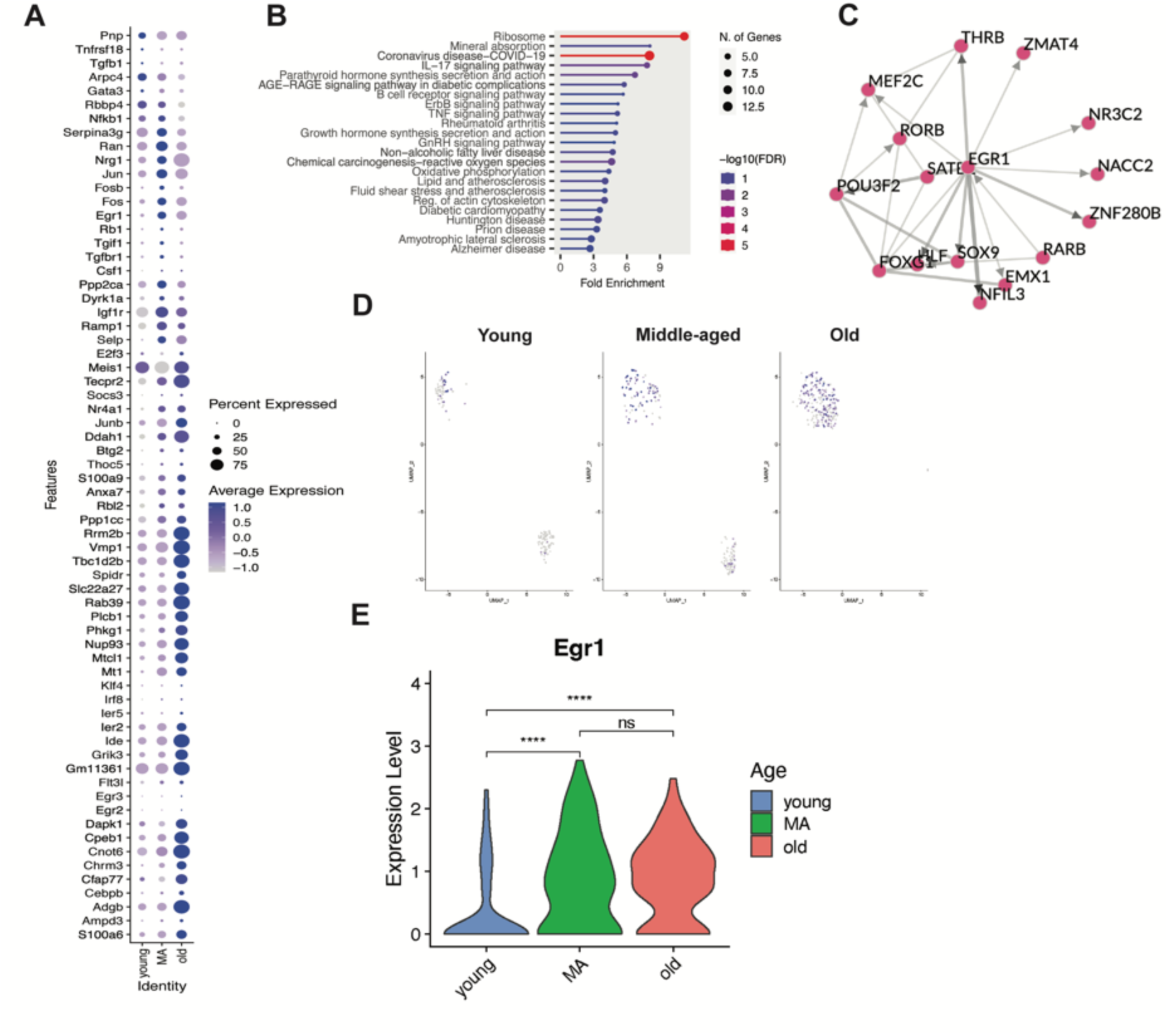
Identifying *Egr1* as a potential master regulator gene in the gene expression signature of aged murine long-term repopulating hematopoietic stem cells. A) Bubble plot of the top downregulated/upregulated gene signature of old LT-HSCs compared to their young and middle-aged counterparts. The color of the spheres indicates the average gene expression, and their size represents the percentage of cells expressing each gene. **B)** The ShinyGO Pathway Analysis^30^ illustrates the top enriched pathways in aged LT-HSCs compared to their young and middle-aged counterparts. The circle size represents the number of differentially expressed genes classified into one specific pathway category. **C)** Transcription factor network derived from the top upregulated genes in aged LT-HSCs based on the ChEA3 analysis ^31^. **D)** UMAP plots showing *Egr1*-expressing cells in young, middle-aged, and old LT-HSC samples. **E)** Violin plots showing *Egr1* expression in young, middle-aged, and old LT-HSC samples; p-values from two-tailed unpaired Student’s t-test, indicating a p-value less than 0.0001 (****) or no significance (ns).

Next, we performed ShinyGO Pathway Analysis^30^ to identify significantly enriched cellular pathways in aged LT-HSCs in an unbiased manner. We utilized the complete list of upregulated genes in old LT-HSCs compared to their middle-aged and young counterparts, respectively. The gene ontology category “ribosome” was the most significantly enriched gene set, which was an expected finding given the known altered upregulation in ribosomal gene transcription with hematopoietic aging, from which others have inferred that old HSPCs may be aberrantly activated through ribosomal biogenesis despite cycling less than younger cells^32^. The rest of the enriched pathways were mainly metabolism-related or linked to the pathogenesis of age-related diseases, such as cardiovascular or degenerative disorders (**Figure 5B**). Using a web-based transcription factor (TF) enrichment analysis tool, ChEA3^31^, we identified EGR1 as the core member of the most probable TF network responsible for the shift in the gene transcription profile of old LT-HSCs (**Figure 5C**).

UMAP analysis revealed that although the expression of *Egr1* was not restricted to middle-aged and old LT-HSCs, its expression level notably increased in middle age, as observed in the bubble plot (**Figure 5A**). Furthermore, *Egr1* was widely expressed within the single LT-HSC cluster seen in older mice. These UMAP plots also showed that while the young and middle-aged groups had the same three clusters, the old LT-HSCs (**Figure 5D**) were absent in the bottom cluster (cluster rep in **Figure 4F**, which had an enriched expression of genes involved in DNA repair and replication). This might be a consequence of the age-related DNA repair defects and subsequent downregulation of genes involved in such pathways or merely an observation derived from a loss of heterogeneity in old LT-HSCs driven by age-related clonal hematopoiesis. Indeed, the expression level of *Egr1* was found to be statistically significant when comparing young versus middle-aged or young versus old LT-HSCs (**Figure 5E**). These results suggest that the upregulation of *Egr1* in middle age might be responsible for a subsequent gene program upregulation promoting murine LT-HSC aging, with widespread *Egr1* constitutive expression in old age to maintain it.

Overall, these data demonstrate that the PURE-seq pipeline can enrich and sequence rare cell populations, such as murine LT-HSCs, to generate high-quality single-cell transcriptomes and, in so doing, give valuable insights into complex biological processes, as it is hematopoietic aging. Compared with existing pipelines, PURE-seq offers a user-friendly solution requiring significantly fewer cells while delivering comparable quality data, which is suitable for biological analyses of rare cell populations.

## Discussion

PURE-seq enables the recovery and sequencing of rare cells from complex cellular populations by integrating two commercially available platforms: FACS and PIP-seq. PIP-seq allows cell barcoding within standard Eppendorf tubes—commonly used vessels for cell recovery in FACS protocols. This direct integration eliminates additional cell transfer steps, significantly reducing cell loss and enabling the reliable capture and sequencing of extremely rare cells.

Our study demonstrates that PURE-seq can enrich and analyze murine LT-HSCs comparably to current methods, such as 10X Genomics, even when using only half of the input cells. This approach is cost-effective, compatible with readily available commercial systems, and opens doors for proteomic analysis, including technologies like CyTOF^33^ and Abseq^34^, as well as multiomics through CITE-seq^35^. PURE-seq has the potential to significantly contribute to genomic and proteomic investigations, particularly those focusing on extremely rare cell populations that can be enriched using flow cytometry. Furthermore, PIP-seq can be combined with antibody-based cell hashing^12^. Although we did not perform hashing in this study, it can be used to further increase the number of cells and conditions processed in the PIP-seq pipeline. In this study, we applied PURE-seq to study hematopoietic aging in murine LT-HSCs. Our results show that LT-HSC heterogeneity is similar in young and middle age but decreases in old mice. We also found that old LT-HSCs exhibit reduced cycling and remain primarily in the G1 phase at the expense of the G2/M and S phases, as previously shown by Hérault *et al*.^21^ Furthermore, our results suggest that EGR1 may be a key TF regulating LT-HSC gene expression during aging, thereby controlling the upregulation of an age-related gene program. Interestingly, *Egr1* expression increases in middle age, potentially indicating its role as an early master regulator of LT-HSC aging, further reinforcing the notion that hematopoietic aging starts in middle age^27^.

While prior studies have shed some light on LT-HSCs^36,37^, the role of *Egr1* in murine LT-HSC aging has not yet been fully elucidated. Recent studies involving scRNA-seq and bulk RNA sequencing have indicated increased *EGR1* expression in aged human HSPCs^15,16^. EGR1 may regulate HSPC quiescence, proliferation, and localization, making it crucial in determining their function and fate. It has been suggested that reducing EGR1 expression may decrease senescence and re-activate aged HSPCs, potentially improving their function and offering a target for hematopoietic rejuvenation strategies^17^. Using PURE-seq, we have identified that *Egr1* may indeed be a master regulator gene of LT-HSC aging in mice, aligning with emerging research in the field and providing a basis for subsequent genomic, epigenomic, and mechanistic studies.

PURE-seq offers significant potential for studying circulating tumor cells (CTCs), which are valuable for research and diagnostics but challenging to capture due to their rarity^38–41^. While positive enrichment using markers like EpCAM, HER2, and MUC1 is common^40,41^, PURE-seq’s throughput enables negative enrichment, allowing it to capture CTCs that may not express these markers. This capability can help discover novel or unexpected CTC types that current methods might miss. With PURE-seq, sufficient CTCs can be captured for meaningful analysis. Using the yield sorting precision mode, we can leverage high-throughput single-cell sequencing downstream of FACS isolation to recover single CTC transcriptomes, even when mixed with non-CTCs. Although this approach may increase false positives, scalable single-cell sequencing can still identify the relevant CTCs, offering a less biased and useful method for diagnostics and monitoring measurable residual disease at low levels.

## Methods

### PURE-seq workflow

PURE-seq combines Fluorescence-activated cell sorting (FACS) and Particle-templated instant partition sequencing (PIP-seq) in an integrated workflow. For the mouse-human mixing experiments described herein, the BD FACS Aria system was used for sorting, and “Sweetspot” was turned on to ensure a stable stream during the sorting. The cooling unit was set to 4°C to keep the collection unit with PIP-seq reaction tube cold throughout the sort. A 0.5 mL tube adapter (Cole-Parmer, EW-17414-73) was inserted into the Aria 1.5 mL collection tube holder to hold the PIP-seq T2 tube. Then, we fine-tuned cell sort stream alignment by using an empty 0.5 mL Eppendorf tube to make sure the test sort droplet was located at the center of the lid when the lid was closed and at the center of the tube bottom when the lid was open. For quality control of each sorting session, we quantified the sorting recovery rate by sorting 100 Calcein labeled cells into a 0.5 mL Eppendorf tube pre-loaded with 10 µL media and counted the number of cells collected under the microscope. The recovery rate is calculated as # Target cells counted under the microscope / # Target cells reported to have been sorted by the instrument. To optimize cell viability and capture efficiency, we capped the total sorting duration to 60 minutes and the total sorted volume to 5 µL (2,500 drops with 85 µM nozzle). Based on BD FACS Aria’s instrument specifications, we limited the flow rate to no more than 8 kHz to minimize shear stress during sheath flow focusing (i.e., 8,000 events per second with 85 µm nozzle). Once the sorting was complete, the PIP-seq T2 tube was unloaded to proceed to the standard PIP-seq protocol from Cell Capture and Lysis after the cell loading step to the preparation of the scRNA-seq library.

### Mouse-human mixing experiment

Human HEK 293T and mouse NIH 3T3 cells (ATCC) were cultured in Dulbecco’s modified Eagle’s medium (DMEM, Thermo Fisher, 11995073) supplemented with 10% fetal bovine serum (FBS; Gibco, 10082147) and 1× Antibiotic-Antimycotic (Gibco, 15240062) at 37°C and 5% CO2. Cells were treated with 0.05% Trypsin-EDTA with Phenol red (Gibco, 25200114) for 3 min, quenched with growth medium, and centrifuged for 3 min at 300*g*. The supernatant was removed, and the cells were resuspended in 1X DPBS without calcium or magnesium. Fresh-frozen human peripheral blood mononuclear cells (PBMCs) were obtained from STEMCELL Technologies. DMEM with 10% FBS was warmed up to 37°C, and the frozen PBMCs were thawed by adding 1 mL of warm media on top of the frozen cells and immediately transferring the media to a 15-mL conical. This process was repeated until all PBMCs were thawed and transferred. Cells were centrifuged for 3 min at 300*g* and resuspended in 1X DPBS. For the 10^-3^, 10^-4^, and 10^-5^ target cell fraction samples, human HEK 293T cells were the target population mixed with mouse NIH 3T3 cells background population. For the 10^-6^ target cell fraction sample, mouse NIH 3T3 cells were the target population mixed with the human PBMCs background population. The target population was treated with 1 μg/mL Calcein Red-Orange (Invitrogen, C34851), and the background population was treated with 1 μg/mL Calcein Green (Invitrogen, C34852) for 15 min at 37°C, followed by washing and dilution to the final concentration in 1× DPBS with 0.1% BSA. The viability and cell concentration were evaluated by an automated cell counter (Bio-Rad, TC20) after adding Trypan Blue (Gibco, 15250061). The mixed cell suspension was filtered through a 40 µm cell strainer (Flowmi, BAH136800040) and processed through the PURE-seq workflow described above to enrich for Calcein Red-Orange labeled cells. For this experiment, we selected the “yield” sorting mode to ensure as many rare cells were sorted, set the flow rate to 8 kHz, and restricted the sorting duration to 60 minutes or if the total sorted volume of 5 µL (2,500 drops with 85 µm nozzle) was reached. In the sequenced libraries, cell transcriptomes were aligned to human or mouse genome to quantify for PURE-seq sensitivity and specificity.

### Sorting precision modes experiment

Calcein Red-Orange labeled human HEK 293T cells and Calcein Green labeled mouse NIH 3T3 cells were mixed at a ratio of 1:1000. The mixed sample volume was controlled at 1mL. Each sample was processed through the PURE-seq workflow described above using “yield” or “single-cell” sorting precision mode until depletion of sample.

### Experimental animals

The study with primary mice was performed in accordance with institutional guidelines established by Memorial Sloan Kettering Cancer Center under the Institutional Animal Care and Use Committee-approved animal protocol (#07-10-016) and the Guide for the Care and Use of Laboratory Animals (National Academy of Sciences 1996). Mice were maintained under specific pathogen-free conditions in a controlled environment that maintained a 12-hour light-dark cycle, and food and water were provided *ad libitum*. The following mice were used: young (2-3 months old), middle-aged (12-14 months old), and old (18-20 months old) female C57BL/6 mice. Young mice were purchased from the Jackson Laboratories and either used when young or aged in-house until middle age. Old mice were obtained from the National Institute of Aging (NIA) and acclimatized for at least 2 weeks at our facility before use. Mice were healthy, had intact immune systems, and had not undergone any prior procedures before euthanasia. For each cohort, 4-6 mice were used to make 2-3 pooled age-matched bone marrow (BM) samples per group prior to sorting.

### Mouse bone marrow harvesting and sample processing for sorting

Mice were humanely euthanized using CO2. BM cells from their limb bones were isolated and resuspended in FACS buffer (PBS + 2% FBS) by centrifugation at 8,000 × g for 1 minute. After removing red blood cells (RBC) with a commercial lysis buffer (BioLegend, 420302), diluted to 1X with distilled water, single-cell suspensions were depleted of hematopoietic cells committed to a specific lineage using a Lineage Cell Depletion Kit (EasySep, StemCell Technologies, Inc., 19856A), according to the manufacturer’s instructions. To label LT-HSC cells, the following fluorophore-conjugated antibodies were used at the indicated dilutions: CD117 (c-Kit) BV785 (clone 2B8, BioLegend; 1:200 dilution), Ly-6A/E (Sca-1) PE/Cy7 (clone D7, BioLegend; 1:1000 dilution), CD48 PerCP/Cy5.5 (clone HM48-1, BioLegend; 1:100 dilution) and CD150 (SLAM) APC (clone TC15-12F12.2, BioLegend; 1:50 dilution). After adding the rat serum and isolation cocktail of the Lineage Cell Depletion Kit, the LT-HSC-labeling antibodies were also added for a 30-minute-long incubation in the dark at 4°C. Following the removal of lineage-positive cells, samples were spun down in FACS buffer and subsequently resuspended in 200-300 μL of FACS buffer containing DAPI at a final concentration of 1 μg/mL. Cells from 2/3 age-matched mice were combined to generate each pool sample, with a total of 2 replicates for the young condition and 3 replicates for the middle-aged and old conditions, respectively (total n=10 mice). Before sorting, we also performed the Rmax method to calculate the maximum recovery of the sample sort and a sorting test with horseradish peroxidase (HRP) using a 0.5 mL collection tube containing a drop of a 3,3’,5,5’-tetramethylbenzidine (TMB), which turned blue if the HRP fell directly into the tube center. Leveraging this HRP-TMB reaction, we ensured that the instrument alignment was correct so that the sample was sorted straight into the PIP-seq T2 reaction. All the mouse primary samples were sorted using a Spectrally Enabled (SE) five-laser BD FACSymphony™ S6, following the protocol described in the “Pure-seq workflow” section and using the “single-cell” sorting precision mode to maximize the purity level.

### scRNA-seq library preparation and sequencing

Single cells were processed for scRNA-seq using the PIP-seq T2 3’ Single Cell RNA kit (v3.0) according to the manufacturer’s protocol (Fluent Biosciences, FB0001026). cDNA and final library DNA quality were confirmed using a 2100 Bioanalyzer Instrument (Agilent Technologies). Libraries were pooled at equimolar ratios and sequenced on an Illumina NovaSeq 6000 S4 platform at PE100 (200 cycles), targeting >50,000 reads per cell. Library demultiplexing, read alignment, identification of empty droplets, and UMI quantification were performed with PIPseeker 1.0.0 (Fluent BioSciences) with default parameters.

### scRNA-seq data analysis in mice

Filtered feature matrices were imported into Seurat, and all downstream analyses were performed using Seurat v4.3.0^42^. For quality control, data were filtered to remove outliers in gene count, UMI count, mitochondrial genes, and ribosomal genes. The 8 samples (young 1-2, middle-aged 1-3, and old 1-3) were normalized by SCTranform and then integrated by Seurat integration using default parameters (SelectIntegrationFeatures and FindIntegrationAnchors), succeeded by normalization and scaling steps^42^. The combined post-sort dataset contained 6,725 cells (Figure), while the pre-sort sample had 40,137 cells. On the complete data, a PCA was estimated, and clustering was performed on 20 principal component dimensions (selected by visual analysis of an Elbowplot) with a resolution of 0.9. A uniform manifold approximation and projection (UMAP) embedding was calculated using the selected 20 principal components as input. Cell cycle was not regressed.

As LT-HSCs were of interest in this study, hematopoietic cells co-expressing the developmental markers *c-Kit*, *Ly6a*, and *Slamf1* were extracted, re-embedded, and re-clustered, followed by a second post-clustering quality control step for further in-depth analysis. From the identified clusters, differential gene expression analysis was conducted using the Seurat function FindAllMarkers to identify genes that were significantly up/downregulated in specific cell clusters compared to others.

After Seurat integration and clustering, different cell types were annotated using the ScType automated cell type classification^43^ with custom markers from the previously published dataset generated by Héuralt *et al.*^21^, where they identified a total of 15 subtypes of LT-HSCs, including 6 primed types (pMast, pNeu, pEr, pL2, pL1, pMk) and 9 non-primed types (div, rep, diff, np4, np3, ifn, np2, np1, tgf). We input the gene markers of these 15 subtypes as a custom marker set to score the cluster markers in our dataset using the ScType R package. Low ScType score clusters (i.e., less than a quarter of the number of cells in a cluster) were considered low-confident and thus designated as “unknown” cell types.

The purity of LT-HSCs in the data was evaluated using the scGate R package^24^. We manually defined a gating model based on the LT-HSC features (*Ptprc* (*CD45*)^+^, c-*Kit*^+^, *Ly6a*^+^, *Slamf1*^+^). The model annotated cells as either “pure” or “impure” based on each cell gene expression. No mouse sample was excluded from these scRNA-seq analyses.

## Data availability

Sequencing data were deposited into the NCBI Gene Expression Omnibus under GSE273803.

## Code availability

The open-source software, tools, and packages used for data analysis in this study, as well as the version of each program, were R (v3.6.1), PIPseeker (v1.0.0), Seurat R package (v4.3.0), scGate R package (v1.6), ScType R package (v1.0), SingleR R package (v1.0). No custom software, tools, or packages were used.

## Acknowledgments

This work was supported by grants R01AI149699 and R01NS130876. I.F-M. was supported by a postgraduate fellowship from the La Caixa Foundation, a Momentum Fellowship from the Mark Foundation for Cancer Research, a Scholarship of Excellence Rafael del Pino, and an NCI F99 award (CA284253-01). R.L.B. was supported by a Damon Runyon-Sohn Fellowship and the NCI (K99CA248460). R.L.L. was supported by a Memorial Sloan Kettering Cancer Center Support Grant/Core Grant P30 CA088748, an R35 grant from the National Institute of Cancer (CA197594), and a collaborative U01 Research Project grant from the National Institute of Aging—the U01 grant was jointly received with the laboratory of Jennifer Trowbridge at the Jackson Laboratories (U01AG077925; 210374-0622-02). We are grateful to members of the Abate laboratory for helpful discussions. We thank Eric Chow and the staff of the UCSF Center for Advanced Technology for their technical support, the members of the Flow Cytometry and the Integrated Genomics Operation (IGO) cores at Memorial Sloan Kettering Cancer Center for their advice and technical help, Kristina Fontanez, Autumn Cholger and Bob Wikle from Fluent BioSciences for their advice and support.

## Contributions

S.P, K.C, A.R.A designed the study; S.P and K.C optimized the PURE-seq workflow; I.F.-M. designed and performed the experiments for all mouse studies; I.F.-M, K.C., S.P analyzed scRNA-seq data; S.V.H. provided bioinformatic and data curation support; M.G.W. assisted with mouse dissections and sample processing; R.L.B. provided input on data visualization; S.P., I.F.-M., A.R.A wrote the manuscript; R.L.L. revised the manuscript; all authors read, reviewed, and approved the manuscript.

## Supplementary Figures

**Figure S1.**
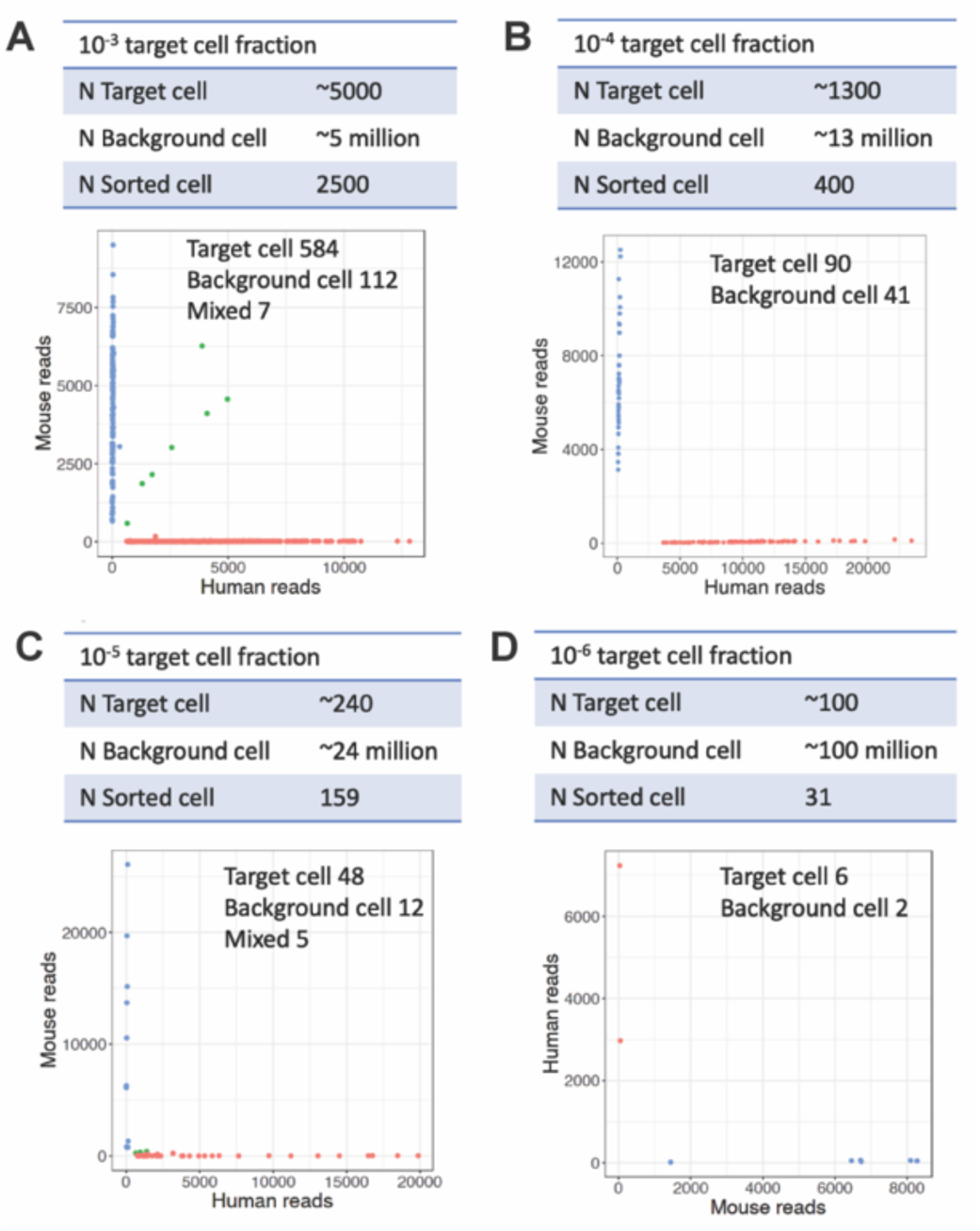
Barnyard plots of 10^-3^, 10^-4^, 10^-5^ and 10^-6^ target cell fractions after sorting. In each table, cell numbers for the corresponding dilution experiment sample are shown (N Target cell and N Background cell) and the number of sorted cells reported by FACS software is noted (N Sorted cell). In each barnyard plot, cells are colored by cell type (blue, mouse reads; red, human reads; green, mixed reads). **A-C)** Human HEK 293T cells and mouse NIH 3T3 cells were stained with Calcein Red-Orange and Calcein Green, respectively. Calcein Red-Orange-positive HEK 293T cells were sorted into PIPseq tubes. **D)** Mouse NIH 3T3 cells and human PBMCs were stained with Calcein Red-Orange and Calcein Green, respectively. Calcein Red-Orange-positive NIH 3T3 cells were sorted out as target cells.

**Figure S2.**
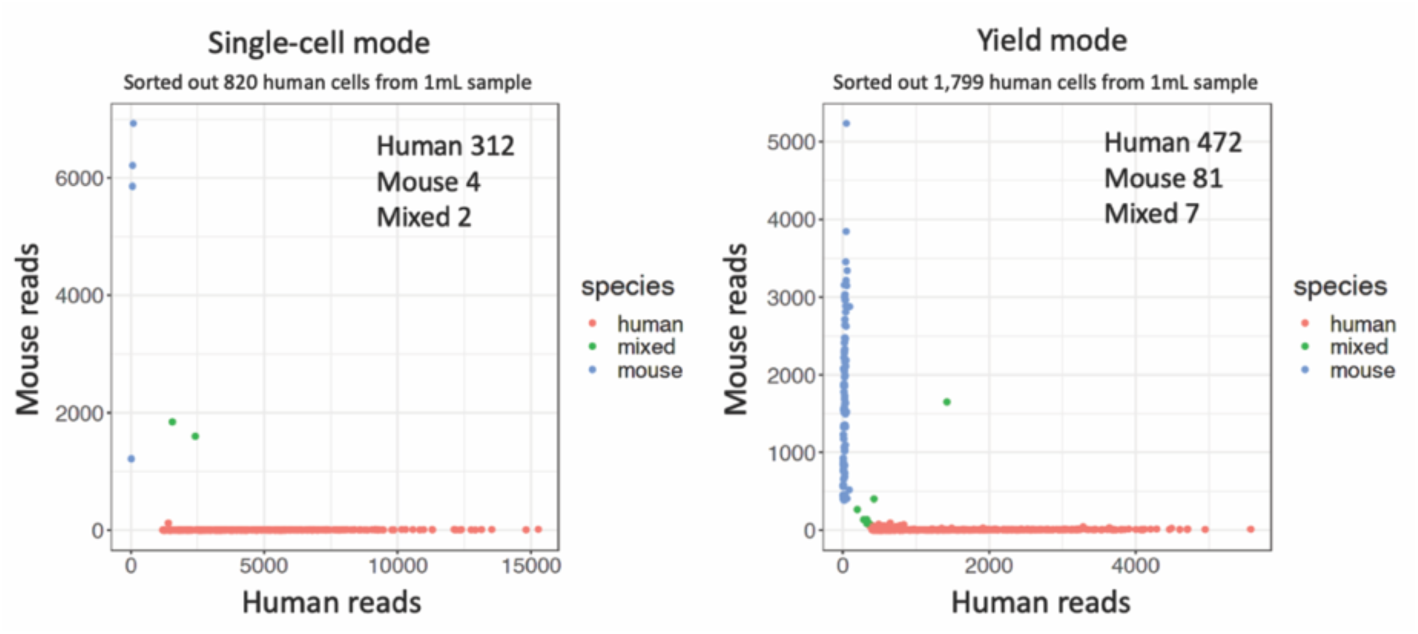
Recovery rate comparison of single-cell and yield sorting precision modes of FACS. In each barnyard plot, cells are colored by cell type (blue, mouse reads; red, human reads; green, mixed reads). Target cell fraction was 10^-3^ and the sample volume was controlled at 1mL. Compared with single-cell mode, yield mode sorted out 2-fold the number of total cells, and sequenced 1.5-fold the number of target rare cells from identical spike-in samples. The purities of single-cell and yield modes were 98% and 84%, respectively.

**Figure S3.**
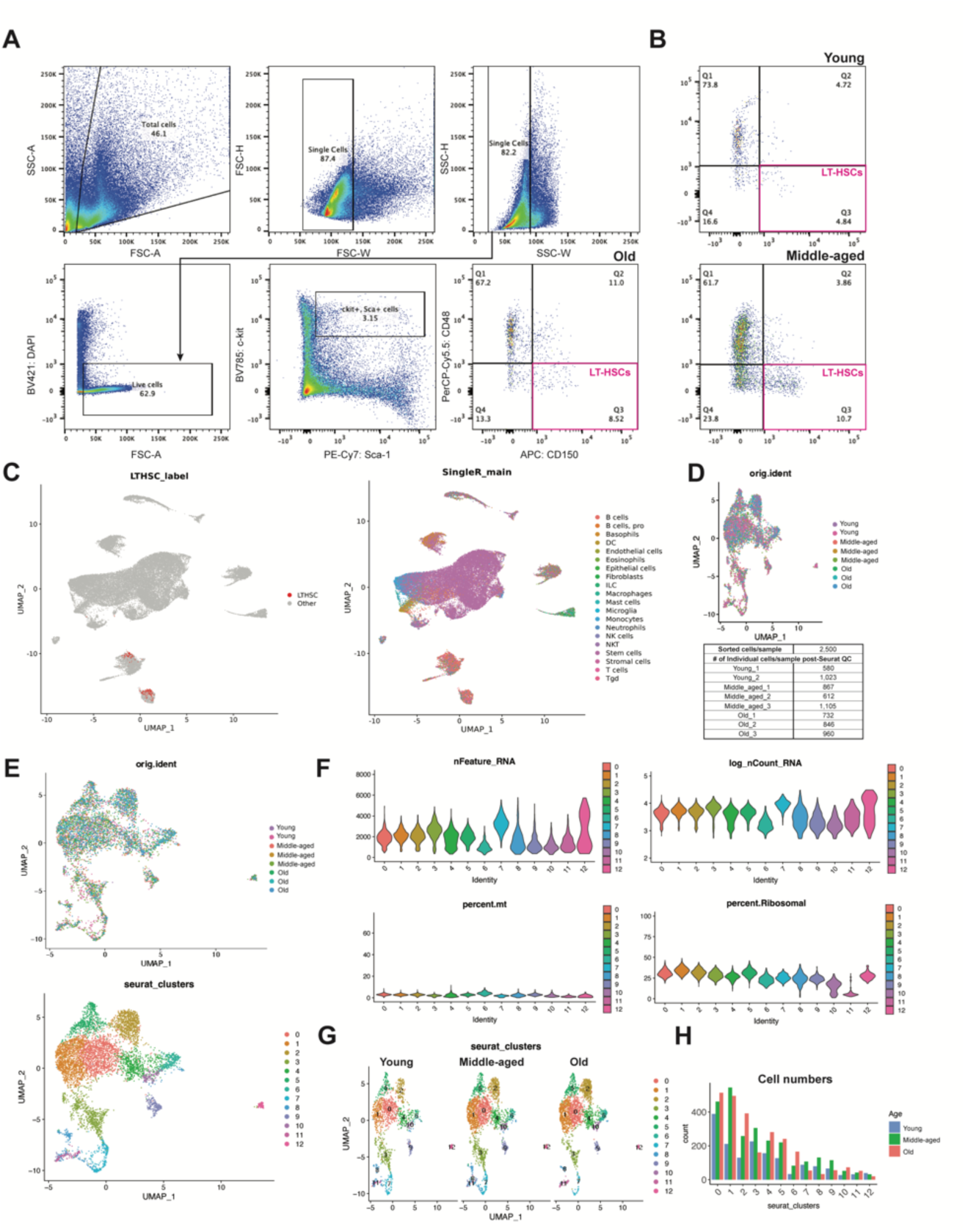
Sorting of murine long-term repopulating hematopoietic stem cell and quality control analysis A) Representative FACS plots using the gating strategy to sort LT- HSCs using old cells as an example. **B)** Representative FACS plots for young (top) and middle-aged (bottom) LT-HSCs. **C)** UMAP plots of pre-sort samples, indicating LT-HSCs as labeled by scGate (left) and unbiased clustering by cell type using the SingleR package^44^ (right). **D)** Integrated UMAP plot of samples from young (n=2), middle-aged (n=3), and old (n=3) mice (top) and the number of sorted cells per sample (n=2,500) and the number of cells recovered after passing quality control standards using the Seurat v4 pipeline, totaling 6,725 cells. **E)** Larger view of the integrated UMAP plot of samples from young (n=2), middle-aged (n=3), and old (n=3) samples, with each age group combining 4-6 mice. Colors indicate the age of the source mice (top) and the clustering of the 6,725 cells using the Seurat v4 pipeline (bottom). **F)** The number of unique genes (nFeature RNA), transcripts (nCount RNA as a logarithmic value), percent mitochondrial reads (percent.mt), and percent ribosomal reads (percent. Ribosomal) as a function of the cluster. **G)** Seurat clustering of young, middle-aged, and old samples. **H)** Bar graph illustrating the cell count for each age group within each Seurat cluster.

**Figure S4.**
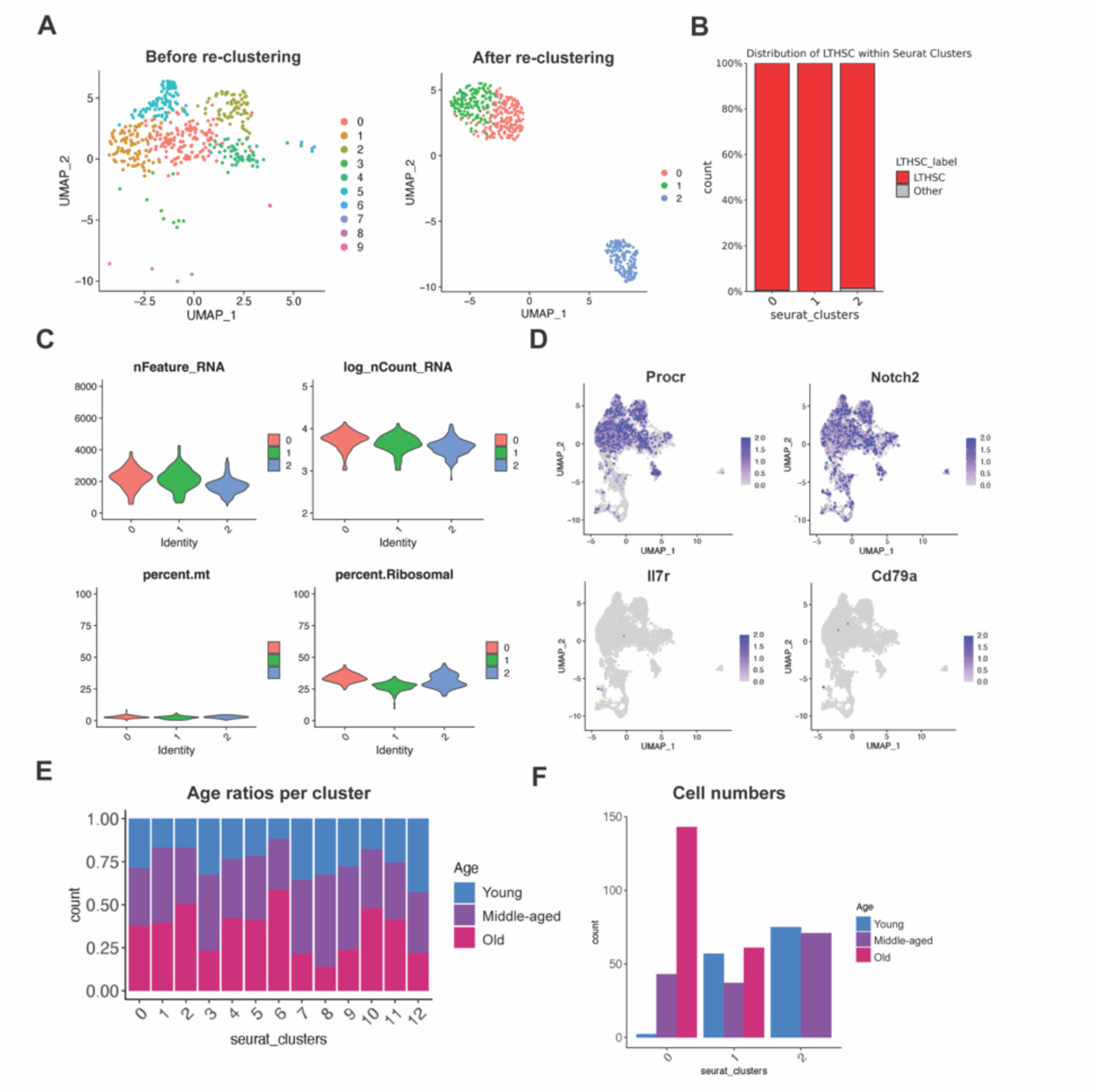
Re-clustering of murine long-term repopulating hematopoietic stem cells, their distribution within Seurat clusters, and quality control post-re-clustering. A) Before (left) and after (right) Seurat re-clustering of purified LT-HSCs according to scGate. **B)** Percentages of LT-HSCs defined by scGate within the Seurat clusters following re-clustering. **C)** The number of unique genes (nFeature RNA), transcripts (nCount RNA as a logarithmic value), percent mitochondrial reads (percent.mt), and percent ribosomal reads (percent. Ribosomal) as a function of the cluster after LT-HSC re-clustering. **D)** UMAP plots colored by expression of selected markers, including undifferentiated HSPC markers (*Procr*, *Notch2*) and markers of lineage bias/commitment (*Il7r*, *Cd79a*). **E-F)** Bar graphs illustrating the cell ratios (left) or counts (right) for each age group within each Seurat cluster subsequent to the re-clustering of LT-HSC.

## Supplementary Tables

**Supplementary Table 1. A**) Cluster marker gene list for integrated dataset after PURE-seq enrichment of LT-HSCs from young (n=2), middle-aged (n=3), and old (n=3) mice samples. **B**) Marker genes for cluster cell-type identification from the Hérault *et al.* dataset.

**Supplementary Table 2.** LT-HSCs identification using scGate analysis for **A**) pre-sort HSC control samples, **B**) PURE-seq enriched LT-HSCs, and **C**) PURE-seq enriched LT-HSCs after reclustering.

## Notes

### Competing Interest Statement

The authors have declared no competing interest.

